# Acriflavine attenuates inflammatory pathology in cutaneous leishmaniasis independently of parasite control

**DOI:** 10.64898/2026.06.05.730355

**Authors:** Erin A. Fowler, Fernanda O. Novais

## Abstract

Cutaneous leishmaniasis is a parasitic skin disease for which current treatments often fail, highlighting the need for new therapeutic approaches. While high parasite burdens are associated with delayed healing, excessive protective immune responses, including elevated IFN-γ production, have also been linked to worse clinical outcomes, indicating that immunopathology contributes significantly to disease progression. Acriflavine is an antimicrobial compound with reported anti-leishmanial activity and the ability to inhibit hypoxia-driven responses. Because *Leishmania*-infected skin is hypoxic in both mice and humans, and hypoxia has been implicated in disease pathogenesis, we investigated whether acriflavine alters the course of cutaneous leishmaniasis by affecting parasite control and/or host immune responses. We found that acriflavine treatment significantly reduced lesion size in *Leishmania major*-infected mice. Unexpectedly, this improvement occurred without changes in parasite burden. Instead, acriflavine treatment reduced the frequency of dendritic cells within lesions and decreased their expression of MHC class II, which correlated with fewer IFN-γ-producing CD4 T cells at the site of infection. These findings indicate that acriflavine ameliorates disease by limiting dendritic cell activation and subsequent IFN-γ-driven immunopathology rather than enhancing parasite clearance. Together, our results identify acriflavine as a potential host-directed therapeutic strategy for cutaneous leishmaniasis and support targeting hypoxia-associated pathways to reduce tissue damage driven by excessive inflammatory responses.

## Introduction

Cutaneous leishmaniasis is a skin disease caused by protozoan parasites of the genus *Leishmania*, characterized by ulcerated skin lesions that can self-heal but may progress to more severe forms if left untreated. In this disease, the immune response plays a dual role by mediating both protective and pathogenic responses that can delay lesion healing despite treatment with anti-parasitic drugs. *Leishmania* parasites infect cells of the innate immune system, primarily macrophages. Control of infection requires antigen presentation by dendritic cells through MHC class II in the presence of IL-12, promoting the differentiation of CD4 T cells into T helper 1 (Th1) cells that produce IFN-γ, which activates infected macrophages to kill *Leishmania* (1). However, while IFN-γ is critical for parasite control, excessive inflammatory responses and sustained IFN-γ production have also been associated with disease exacerbation in some forms of cutaneous leishmaniasis (2–13).

Presently, there is no approved human vaccine for leishmaniasis, and current treatments are limited by toxicity, high failure rates, prolonged treatment regimens, and the emergence of drug resistance (14). Therefore, there is a need to identify alternative therapeutic strategies for cutaneous leishmaniasis. Acriflavine is an acridine-derived compound that has historically been used as an antiseptic and exhibits antimicrobial activity against a broad range of pathogens, including protozoan parasites. Previous studies demonstrated that acriflavine can impair the growth of several protozoan species, including *Leishmania tarentolae*, in part through inhibition of DNA replication (15–19). In addition, treatment with acriflavine in combination with anti-parasitic drugs reduced lesion development and parasite burden in the livers and spleens of *Leishmania major*-infected mice through mechanisms that remain poorly understood (20). Together, these findings suggest that acriflavine may have therapeutic potential in cutaneous leishmaniasis through direct anti-parasitic effects. However, acriflavine has also been shown to modulate host responses.

Recent studies have expanded interest in acriflavine beyond its antimicrobial activity because of its potential to block hypoxia-mediated pathology (21). Hypoxia, or low oxygen tension, is a common feature of many diseases and has been associated with poor prognosis in cancer, neurodegenerative disorders, and chronic inflammatory diseases (22–26). Cellular adaptation to hypoxia is largely mediated by hypoxia-inducible factors (HIFs), transcription factors that regulate the expression of genes involved in inflammation, metabolism, angiogenesis, and cell survival. Under hypoxic conditions, HIF-α subunits become stabilized and promote transcription of hypoxia-responsive genes (23, 27). Among compounds targeting hypoxia signaling pathways, acriflavine has been shown to inhibit the expression of HIF target genes (28–33). Importantly, acriflavine treatment reduced pathology in multiple experimental models, including cancer, metastasis, preeclampsia, and optic neuritis, in which disease progression has been linked to HIF stabilization (28, 29, 31–35). These findings raise the possibility that acriflavine may also modulate inflammatory responses in diseases characterized by tissue hypoxia.

Skin lesions from *Leishmania*-infected mice and patients are hypoxic and exhibit stabilization of HIF-1α, while lesion severity inversely correlates with tissue oxygenation (36–43). Ongoing studies have begun to define the role of hypoxia and HIF signaling in the pathogenesis of cutaneous leishmaniasis, and our group previously demonstrated that lesional hypoxia promotes pathogenic responses by cytotoxic CD8 T cells (40–50). Despite the promising therapeutic potential of acriflavine, little is known about how this compound modulates immune responses during infection. Therefore, we investigated whether acriflavine treatment could reduce lesion development in *L. major*-infected mice by exerting potential antiparasitic and/or immunomodulatory effects.

Here, we show that acriflavine treatment significantly reduced lesion development in *L. major*-infected mice. Unexpectedly, this reduction in pathology occurred independently of parasite control. Instead, acriflavine treatment reduced both the frequency of lesional dendritic cells and their expression of MHC class II, which correlated with a decrease in the number of IFN-γ-producing CD4 T cells within lesions. Together, these findings suggest that acriflavine limits lesion development by modulating local inflammatory responses rather than enhancing anti-parasitic immunity.

## Results

### Acriflavine reduces lesion development independently of parasite control in cutaneous leishmaniasis

To determine whether acriflavine alters the course of cutaneous leishmaniasis, C57BL/6 mice were intradermally infected in the ear with *L. major* and treated daily with either acriflavine (5 mg/kg) or vehicle beginning at the time of infection (Figure 1A). Lesion development was monitored weekly by measuring ear thickness. Acriflavine-treated mice developed smaller lesions beginning two weeks post-infection compared to vehicle-treated controls (Figure 1B), and the area under the curve indicated that disease was significantly decreased in acriflavine-treated compared to vehicle-treated mice (Figure 1C). Given the anti-parasitic properties of acriflavine, we next assessed whether reduced lesion development correlated with better parasite control. Unexpectedly, parasite burden within lesions was comparable between vehicle- and acriflavine-treated mice at 2, 3, 4, and 9 weeks post-infection (Figure 1D). These findings indicate that acriflavine reduces lesion pathology independently of parasite clearance, suggesting that treatment instead may decrease the inflammatory response within lesions.

**Figure 1.**
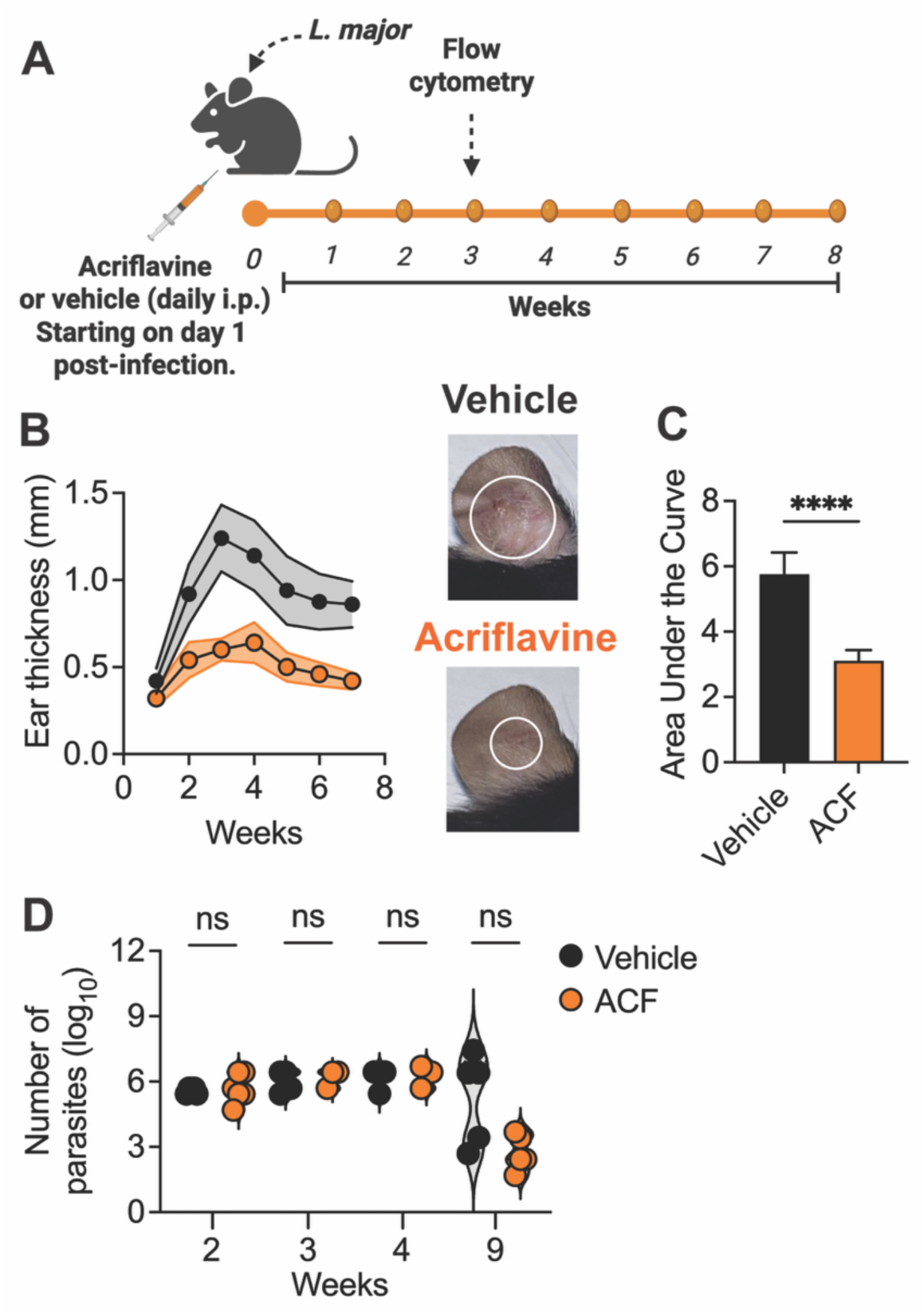
Systemic treatment of *Leishmania*-infected mice with acriflavine reduces skin pathology. (A) Schematic representation of C57BL/6 mice infected with *Leishmania major* and treated daily with intraperitoneal injections of acriflavine or vehicle until 9 weeks post infection. (B) Ear thickness was measured weekly, and representative images of lesions at 3 weeks post-infection are shown. Ear thickness measurement is representative of more than 3 experiments with at least 3 mice per experimental group. (C) Area under the curve for ear thickness. (D) Parasite numbers at 2, 3, 4, and 9 weeks of infection. ****P ≤ 0.0001 by 2-tailed Student’s t test (C and D).

### Acriflavine selectively reduces lesional dendritic cells and MHC-II^+^ dendritic cells

To investigate whether acriflavine treatment altered immune responses during infection, we analyzed lesional immune populations at 3 weeks post-infection (Supplemental Figure 1), the time point at which the greatest difference in lesion size was observed between treatment groups. Flow cytometric analysis revealed no differences in the frequency of total CD45^+^ leukocytes, neutrophils, macrophages, or monocytes between vehicle- and acriflavine-treated mice (Figure 2A-C). In contrast, acriflavine-treated mice exhibited a significant reduction in the frequency of lesional dendritic cells (Figure 2D).

**Figure 2.**
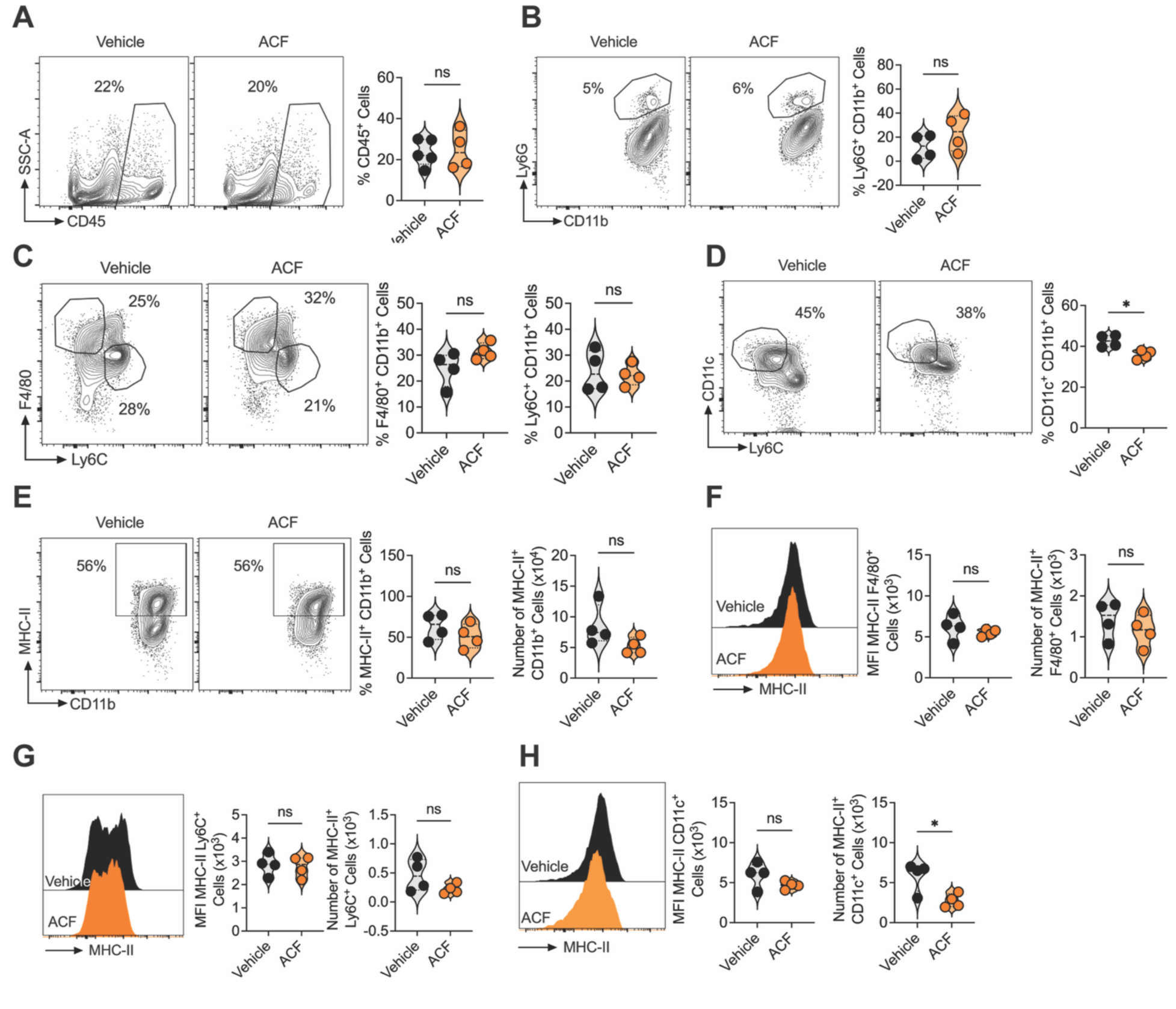
Treatment of *Leishmania*-infected mice with acriflavine reduces the number of MHC-II^+^ dendritic cells. C57BL/6 mice were infected with *Leishmania major* and treated daily with intraperitoneal injections of acriflavine or vehicle for 3 weeks, and single-cell suspensions of lesions and draining lymph nodes (dLN) were analyzed by flow cytometry. The frequency of (A) CD45^+^, (B) neutrophils, (C) macrophages and monocytes, (D) dendritic cells, and (E) MHC-II^+^ CD11b cells in the lesion was assessed by flow cytometry. (left) Mean fluorescent intensity (MFI) and (right) number of MHC-II expression on (F) macrophages, (G) monocytes, and (H) dendritic cells in the lesion. (A-H) Representative contour plots, histograms, and scatter dot plots showing individual mice are representative of 2 experiments with at least 4 mice per experimental group. *P ≤ 0.05 by 2-tailed Student’s t test (A-H). Gating strategy: (A) live, single cells, CD45; (B) live, single cells, CD45, CD11b, Ly6G; (C, D, F, G, and H) live, single cells, CD45, CD11b, Ly6G^-^, and F4/80 or Ly6C or CD11c; (E) live, single cells, CD45, CD11b.

Because dendritic cells are critical antigen-presenting cells during *Leishmania* infection, we next evaluated MHC class II expression within lesional myeloid populations. Acriflavine treatment did not alter the frequency or number of MHC-II^+^ myeloid cells (CD11b^+^) within lesions (Figure 2E). Similarly, MHC-II expression on a per-cell basis, as assessed by mean fluorescence intensity (MFI), was not significantly altered in macrophages, monocytes, or dendritic cells (Figure 2F-2H). However, acriflavine treatment resulted in an approximately two-fold reduction in the number of MHC-II^+^ dendritic cells within lesions (Figure 2H), while other myeloid cells were not impacted (Figure 2F-G). Together, these findings indicate that acriflavine treatment selectively reduces lesional dendritic cells and MHC-II expression rather than broadly suppressing myeloid cell accumulation or MHC-II expression.

### Acriflavine treatment reduces the accumulation of lesional T cells

Because acriflavine treatment reduced dendritic cells and MHC-II expression within lesions, we next investigated whether T cell subsets were also altered during infection. T cell responses were analyzed 3 weeks post-infection (Supplemental Figure 2), the time point with the greatest difference in lesion size between treatment groups. Acriflavine-treated mice exhibited a reduction in the frequency of total T cells within both the draining lymph node (dLN) and lesions compared to vehicle-treated controls. This reduction corresponded to only a decrease in the number of lesional T cells in acriflavine-treated mice (Figure 3A). We found no significant differences in the frequency of antigen-experienced (CD44^high^) CD4 T cells in dLN or lesion, but there was a reduction in the number of lesional CD4 T cells in acriflavine-treated compared to vehicle-treated mice (Figure 3B). Additionally, acriflavine treatment significantly reduced the frequency and number of lesional CD44^high^ CD8 T cells, while no changes were observed in the dLN (Figure 3C).

**Figure 3.**
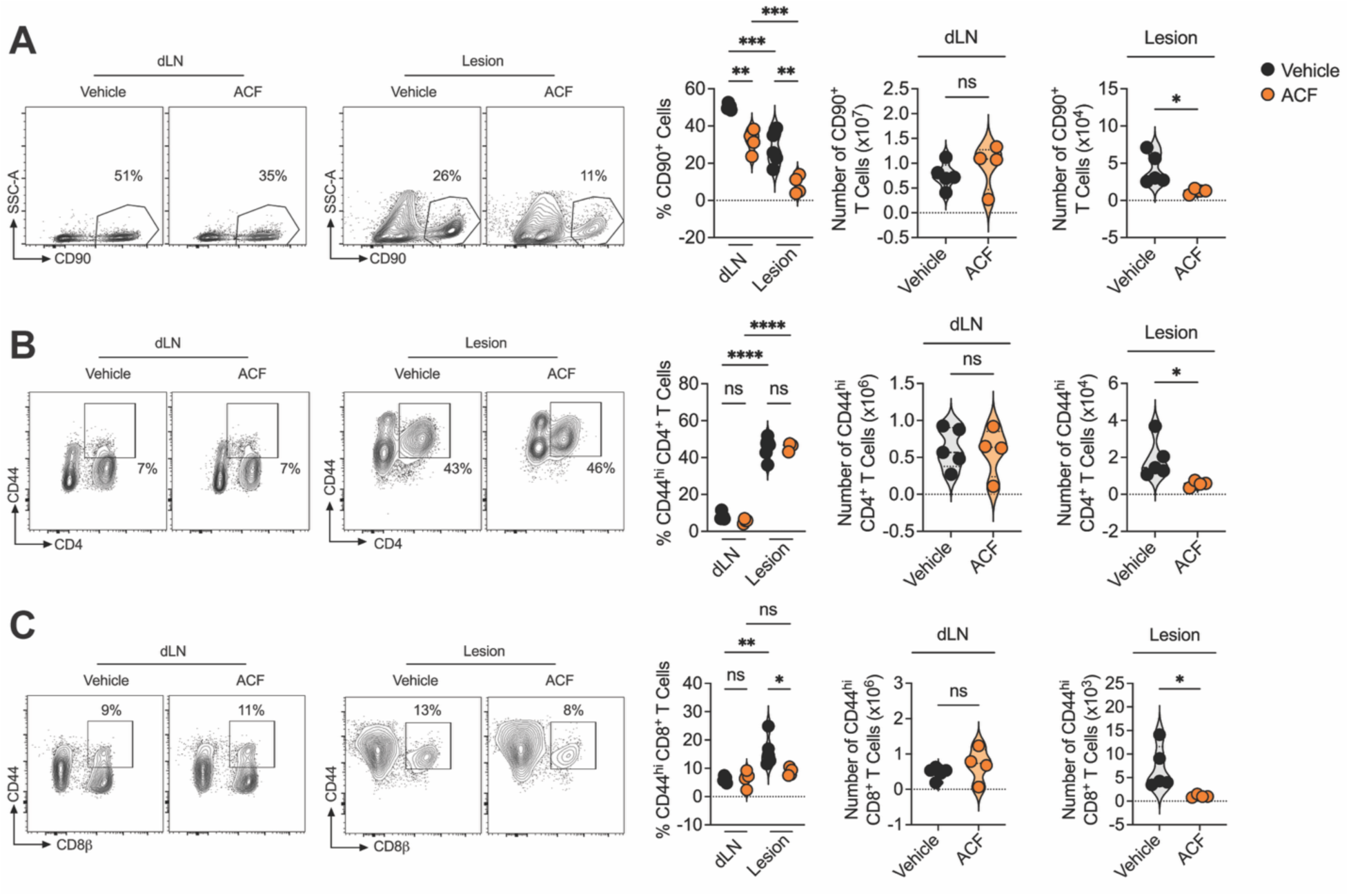
Treatment of *Leishmania*-infected mice with acriflavine reduces the number of T cells in the lesion. C57BL/6 mice were infected with *Leishmania major* and treated daily with intraperitoneal injections of acriflavine or vehicle for 3 weeks, and single-cell suspensions of lesions and draining lymph nodes (dLN) were analyzed by flow cytometry. The frequency and total number of (A) CD90^+^ T cells, (B) CD44^hi^ CD4^+^ T cells, and (C) CD44^hi^ CD8^+^ T cells in the dLN and lesions. (A-C) Representative contour plots and scatter dot plots showing individual mice are representative of more than 3 experiments with at least 3 mice per experimental group. (A-C) *P ≤ 0.05, **P ≤ .01, ***P ≤ 0.001, and ****P ≤ 0.0001, by 2-tailed Student’s t test (number) and 1-way ANOVA (frequency). Gating strategy: (A) live, single cells, CD45, CD90; (B-D) live, single cells, CD45, CD90, CD8 or CD4, CD44^high^.

Because regulatory T cells (Tregs) can modulate inflammatory responses during cutaneous leishmaniasis (51–56), we also evaluated this population following treatment. Acriflavine treatment did not alter the frequency or number of CD44^high^ FoxP3^+^ Tregs within either the dLN or lesions (Supplemental Figure 3). Together, these findings indicate that acriflavine treatment selectively reduces effector T cell accumulation within lesions.

### Acriflavine treatment reduces the number of IFN-γ-producing T cells within lesions

Given the reduction in dendritic cells and lesional T cells following acriflavine treatment, we next assessed whether T cell activation and effector responses were altered during infection. Programmed cell death protein-1 (PD-1) is an inhibitory receptor induced on T cells following activation through the T cell receptor and is further upregulated by persistent antigen stimulation and inflammatory signals. Engagement of PD-1 by its ligands dampens T cell proliferation, cytokine production, and effector function, serving as a critical mechanism to limit immunopathology and maintain immune homeostasis (57). PD-1 expression was elevated on lesional CD44^high^ CD4 and CD8 T cells compared to cells within the dLN (Figure 4A-B), consistent with previous reports (40, 58–61). Acriflavine treatment reduced the number of PD-1^+^ CD4 T cells (Figure 4A) and the frequency of PD-1^+^ CD8 T cells (Figure 4B) in lesions, suggesting altered activation of lesional T cells upon acriflavine administration.

**Figure 4.**
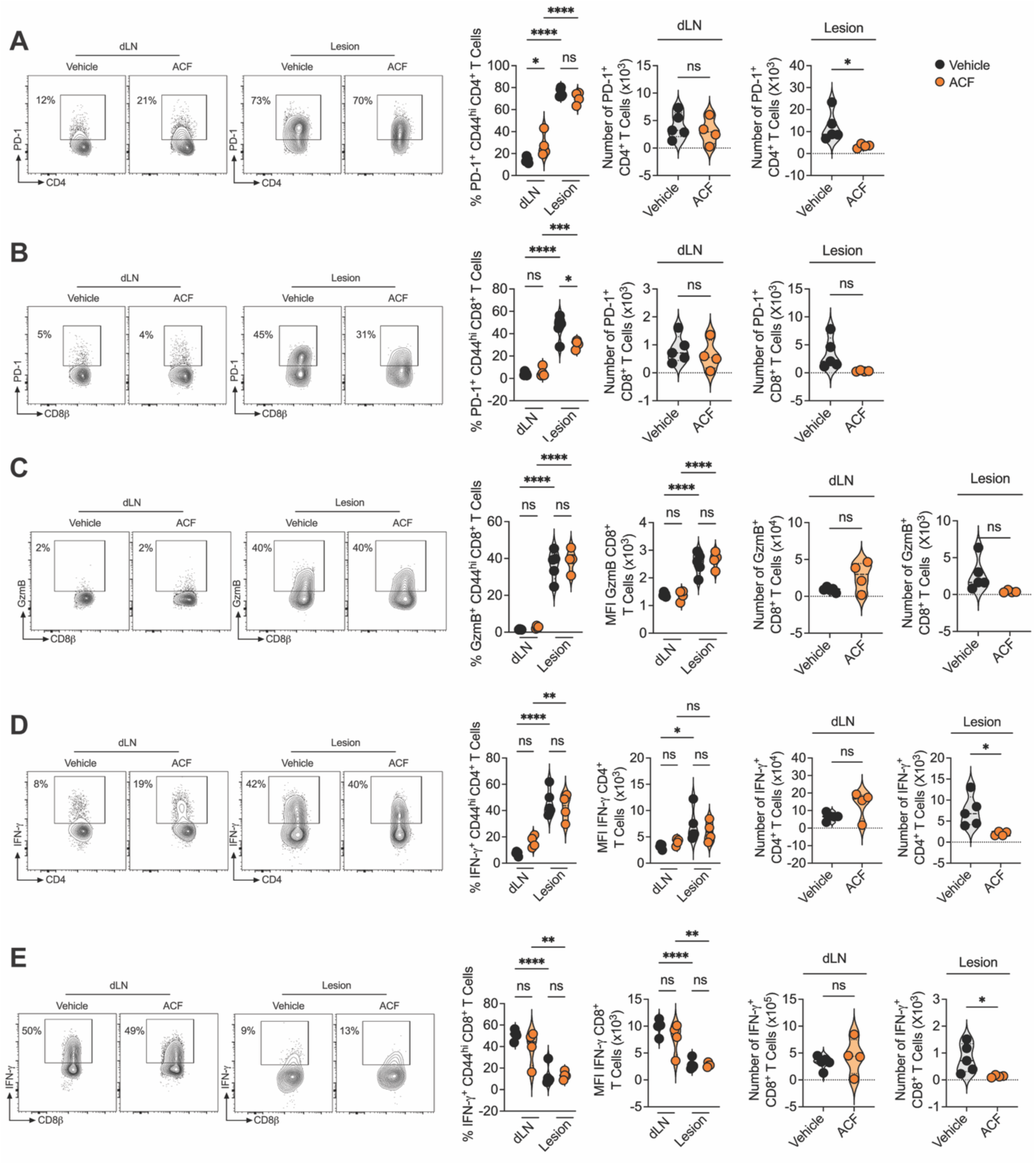
Treatment of *Leishmania*-infected mice with acriflavine reduces the number of IFN-γ^+^ T cells. C57BL/6 mice were infected with *Leishmania major* and treated daily with intraperitoneal injections of acriflavine or vehicle for 3 weeks, and single-cell suspensions of lesions and draining lymph nodes (dLN) were analyzed by flow cytometry. The frequency and number of PD-1^+^ (A) CD4 and (B) CD8 T cells in the dLN and lesions. (C) The frequency, mean fluorescent intensity (MFI), and number of granzyme B^+^ (GzmB) CD8 T cells in the dLN and lesions. (D-E) The frequency, MFI, and number of IFN-γ^+^ (D) CD4 and (E) CD8 T cells in the dLN and lesions. Representative contour plots, histograms, and scatter dot plots showing individual mice are representative of (A, B, D, and E) 2 experiments with at least 4 mice per experimental group or (C) more than 3 experiments with at least 3 mice per experimental group. *P ≤ 0.05, **P ≤ .01, ***P ≤ 0.001, and ****P ≤ 0.0001, by 2-tailed Student’s t test (number) and 1-way ANOVA (frequency and MFI). Gating strategy: (A and D) live, single cells, CD45, CD90, CD4, CD44^high^, FoxP3^-^ (B, C, and E) live, single cells, CD45, CD90, CD8, CD44^high^.

We next assessed whether acriflavine altered expression of the cytotoxic molecule granzyme B, which has been implicated in CD8 T cell-mediated pathology during cutaneous leishmaniasis. Acriflavine treatment did not significantly alter the frequency, MFI, or number of granzyme B^+^ CD8 T cells in lesions of dLN (Figure 4C).

Because excessive IFN-γ production contributes to pathology in cutaneous leishmaniasis, we next evaluated IFN-γ expression by T cells in the dLN and lesions of infected mice following acriflavine treatment, compared with vehicle-treated mice. While acriflavine treatment did not alter the frequency or MFI of IFN-γ within CD4 or CD8 T cells in either the dLN or lesions, treated mice exhibited reduced numbers of IFN-γ^+^ CD4 and CD8 T cells within lesions (Figure 4D-E). Together, these findings indicate that acriflavine treatment primarily alters the accumulation of effector T cells, particularly those expressing IFN-γ within lesions rather than broadly suppressing T cell effector function.

## Discussion

Acriflavine has shown preclinical promise as both an antimicrobial and a therapeutic agent capable of limiting hypoxia-associated pathology in several disease settings (17–21, 28–35, 62–64). Here, we demonstrate that acriflavine treatment significantly reduced lesion development during experimental cutaneous leishmaniasis independently of parasite burden. Although acriflavine has demonstrated anti-protozoan activity in vitro and in vivo against several pathogens, including *Plasmodium* species, we found no evidence that acriflavine enhanced *L. major* control within lesions (15–19). One previous study investigating acriflavine treatment during *L. major* infection was found to reduce lesion development compared to untreated mice, which was associated with a decrease in the number of parasites in the spleen and liver. No parasite numbers were reported at the site of infection (20). Our findings suggest that the beneficial effects of acriflavine in cutaneous leishmaniasis lesions are attributable to modulation of host inflammatory responses rather than to direct antiparasitic activity.

Despite the promising indication that acriflavine reduces pathology in a variety of hypoxic disease states, there has been minimal investigation into the overall impact of acriflavine on the overall immune response (31, 32, 34, 65). Previous studies from our group established that hypoxia promotes pathogenic responses by cytotoxic CD8 T cells during cutaneous leishmaniasis (40, 44). Therefore, we initially hypothesized that acriflavine treatment would reduce cytotoxic T cell function within lesions. Surprisingly, acriflavine treatment did not substantially alter granzyme B expression by lesional CD8 T cells, despite reducing overall T cell accumulation within lesions. These data indicate that acriflavine treatment altered lesion development independent of cytotoxicity. Instead, our data show a selective reduction in lesional dendritic cells and their expression of MHC-II following acriflavine treatment, despite minimal changes in other myeloid cells. Reductions in dendritic cells following acriflavine treatment have previously been reported in murine models of breast cancer (34), although the impact of acriflavine on dendritic cells beyond accumulation was not investigated. Dendritic cells are critical for sustaining local T cell responses during *Leishmania* infection, and the reduction in dendritic cells observed here correlated with fewer lesional T cells and reduced numbers of IFN-γ-producing T cells. In line, one group showed that acriflavine reduced the accumulation of microglia/macrophages and T cells and subsequent pathology of the optic nerve in the experimental autoimmune encephalomyelitis (EAE) mouse model (32). The authors did not analyze expression of MHC-II in myeloid cells or the effector function of T cells, but disease progression in EAE and multiple sclerosis is strongly linked to excessive IFN-γ and hypoxia (66–72). Together, these results indicate a potential broad application of acriflavine as an immunomodulator of T cell function in hypoxic environments and should be further investigated in relevant disease models such as systemic lupus erythematosus and rheumatoid arthritis (73–77).

The mechanisms through which acriflavine alters immune responses during infection remain incompletely understood. Acriflavine has been shown to inhibit the transcription of hypoxia-inducible factors (HIF) target genes, a major pathway by which cells adapt to a hypoxic environment (28–33). Anders et al. found that acriflavine reduced the proliferation and expression of HIF target genes by splenocytes from EAE mice in a dose-dependent manner. Additionally, work in a mouse model of lupus nephritis showed that knockdown of HIF in renal T cells reduced their survival by altering the splicing of the survival factor BNIP3 (73). However, acriflavine has pleiotropic effects that may independently influence immune cell activation, survival, metabolism, or trafficking (21). Therefore, although our findings are consistent with modulation of hypoxia-associated inflammatory pathways, additional studies will be required to determine the precise mechanisms by which acriflavine alters dendritic cell and T cell responses during infection.

Within the context of cutaneous leishmaniasis, our studies support a large body of evidence demonstrating that excessive inflammatory responses, including sustained IFN-γ production, contribute to tissue damage. Despite the requirement of IFN-γ for *Leishmania* control, there is a paradoxical observation that more severe lesions contain high levels of IFN-γ (2, 3, 6, 7, 9, 11–13). This is most evident in the mucocutaneous form of this disease, where parasites metastasize from the skin to the nasopharyngeal mucosa, a severe disease manifestation that develops in a subset of patients (4, 5, 8, 10). Additionally, transcriptional analysis from our group showed that lesional expression of *IFNG* was increased in patients who failed treatment, and the combined expression of *IFNG, IL1B,* and *GNLY* predicted treatment failure with 86-96% accuracy (12). Polymorphisms in the *IFNG* gene have been linked to cutaneous leishmaniasis susceptibility, with discrepancies likely owing to differences in cohort location and parasite species (78–80). Therefore, it is evident that there is a precarious balance of IFN-γ that must be maintained to preserve parasite killing while limiting skin damage.

Current therapies for cutaneous leishmaniasis primarily target the parasite but do not directly address the immunopathology that contributes to tissue damage and delayed healing (14). Our findings support the concept that host-directed therapies capable of selectively reducing inflammatory pathology may provide therapeutic benefit even without substantially enhancing parasite clearance. Given that acriflavine is inexpensive, clinically accessible, and relatively well tolerated, these findings support further investigation of acriflavine and related compounds as immunomodulatory therapies for cutaneous leishmaniasis and other diseases characterized by hypoxia-associated inflammatory pathology. Importantly, acriflavine treatment did not affect parasite burden, suggesting that the reduction in IFN-γ lessened lesion size while remaining sufficient to control *Leishmania*, indicating that it is a viable therapy that could be used to treat patients with cutaneous leishmaniasis by modulating local immune responses.

## Materials & Methods

### Mice

Our animal experiments were performed using male and female mice, and controls were sex matched, but sex was not specifically tested as a biological variable. C57BL/6 mice (6 weeks) were purchased from Charles River Laboratories. All mice were maintained in a specific pathogen-free environment at The Ohio State University Animal Care Facilities. This study was carried out per the recommendations in the Guide for the Care and Use of Laboratory Animals of the National Institutes of Health. The Institutional Animal Care and Use Committee and The Ohio State University approved the protocol.

### Parasites

*Leishmania major* (strain WHO/MHOM/IL/80/Friedlin) were grown in Schneider’s insect medium (Gibco) supplemented with 20% heat-inactivated fetal bovine serum (Sigma-Aldrich) and 2 mM glutamine (Thermo Fisher Scientific). Metacyclic-enriched promastigotes were used for infection (81). Mice were infected with 10^6^ *L. major* intradermally in the ear, and the lesion progression was monitored weekly by measuring the ear thickness with a digital caliper.

### Parasite titration

The parasite burden in the ears was quantified as described previously (82). Briefly, the homogenate was serially diluted and incubated at 26°C. The number of viable parasites was calculated from the highest dilution at which parasites were observed after 7 days.

### In vivo drug treatment

Acriflavine (Sigma-Aldrich, catalog A8251) or vehicle (PBS) was administered intraperitoneally (5 mg/kg in 200 uL) daily for up to 9 weeks before euthanasia beginning one day after infection.

### Single-cell suspension preparation

Infected ears were collected, the dorsal and ventral layers of the ear were separated, and the ears were incubated in RPMI 1640 (Gibco) with 250 μg/ml of Liberase TL (Roche Diagnostics) and DNAse I (Sigma-Aldrich) and dissociated using the gentleMACS Dissociator with Heaters (Miltenyi Biotec) using program 37_Multi_H then a cell strainer (40 μm, BD Pharmingen), and an aliquot of the cell suspension was used for parasite titration. dLNs were homogenized using a cell strainer (40 μm, BD Pharmingen) to obtain single-cell suspensions.

### Flow cytometric analysis

Before surface and intracellular staining, cell suspensions were stained with LIVE/DEAD fixable blue dead cell stain kit (L23105) according to the manufacturer’s instructions. Cells analysis was performed using the FlowJo Software (Tree Star), and gates were created on the basis of fluorescence minus one control. The following antibodies were used: MHC-II (clone M5/114.15.2, catalog 56-5321-80), CD45 (clone 30-F11, catalog 127614), F4/80 (clone B8, catalog 25-4801-82), FoxP3 (clone FJK-16s, catalog 56-5773-82), Granzyme B (clone GB11, catalog GRB05), Arginase 1 (clone A1exF5, catalog 48-3697-82), IFN-γ (clone XMG1.2, catalog 25-7311-82), CD4 (clone RM4-5, catalog 11-0042-82) (all from Invitrogen, Thermo Fisher Scientific); PD-L2 (clone MIH37, catalog 752608), Ly6C (clone AL-21, catalog 560525), CD44 (clone IM7, catalog 751414) (all from BD Biosciences); CD45 (clone 30-F11, catalog 103137), CD8β (clone YTS156.7.7, catalog 126633), CD90 (clone 53-2.1, catalog 140306), Ly6G: (clone 1A8, catalog 127614), CD11c (clone N418, catalog 117331), CD11b (clone M1/70, catalog 101259), PDL1 (clone 10F.9G2, catalog 124319), iNOS (clone W16030C, catalog 696806), PD1 (clone 29F.1A12, catalog 135224) (all from BioLegend). The stained cells were acquired on Cytek Aurora 4 Laser (UV-V-B-R) or Cytek Aurora 5 Laser (UV-V-B-YG-R).

## Statistics

For differences between 2 groups, statistical significance was determined using an unpaired Student’s t-test. For multiple comparisons, 1-way ANOVA was performed, followed by Tukey’s post hoc test to determine differences between groups. For Figure 1B differences in lesion development were determined using area under the curve. Differences were considered significant when ∗P ≤ .05, ∗∗P ≤ .01, ∗∗∗P ≤ .001, or ∗∗∗∗P ≤ .0001.

## Acknowledgements

This work was supported by National Institutes of Health grant R01AI162711 (to FON), the Host Defense and Microbial Biology Program from the Infectious Diseases Institute at the Ohio State University (to FON), and National Institute of Health training program T32 AI165391“Interdisciplinary Program in Microbe-Host Biology” (to EAF).

## Author Contributions

Conceptualization: FON; Funding Acquisition: FON; Investigation: EAF, FON; Supervision: FON; Writing – Original Draft Preparation: EAF. Writing – Review and Editing: FON

---

**Supplemental Figure 1.**
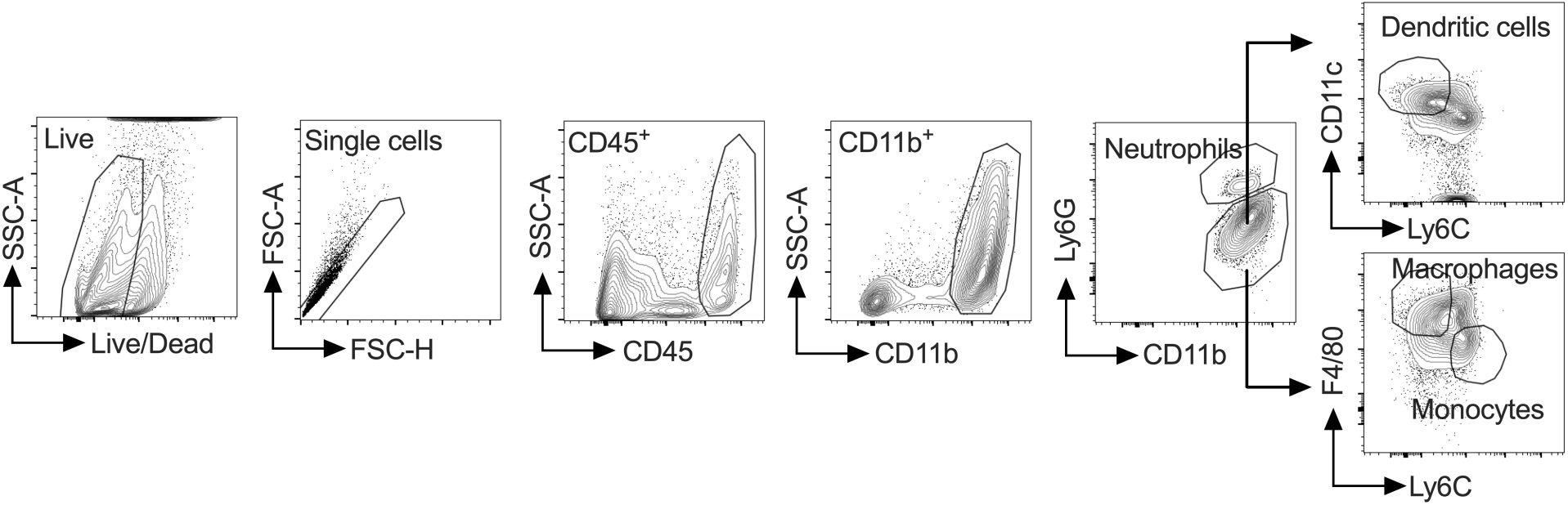
Schematic representation of the gating strategy for myeloid populations for Figure 2.

**Supplemental Figure 2.**
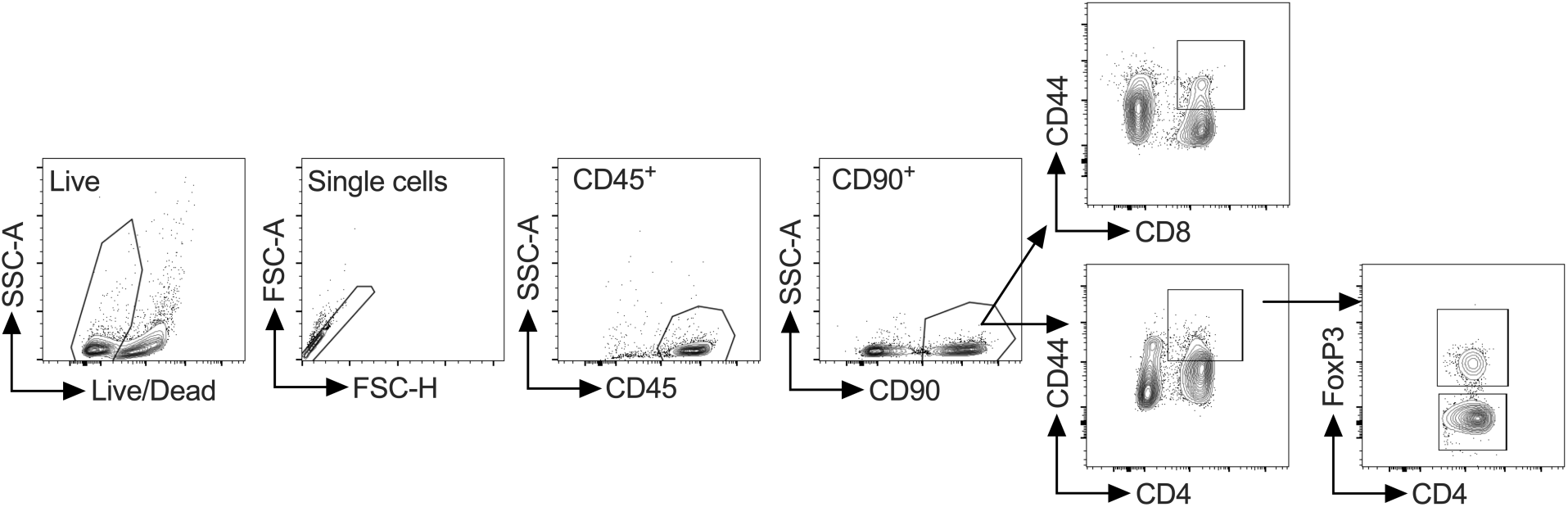
Schematic representation of the gating strategy for T cell populations for Figures 3 and 4.

**Supplemental Figure 3.**
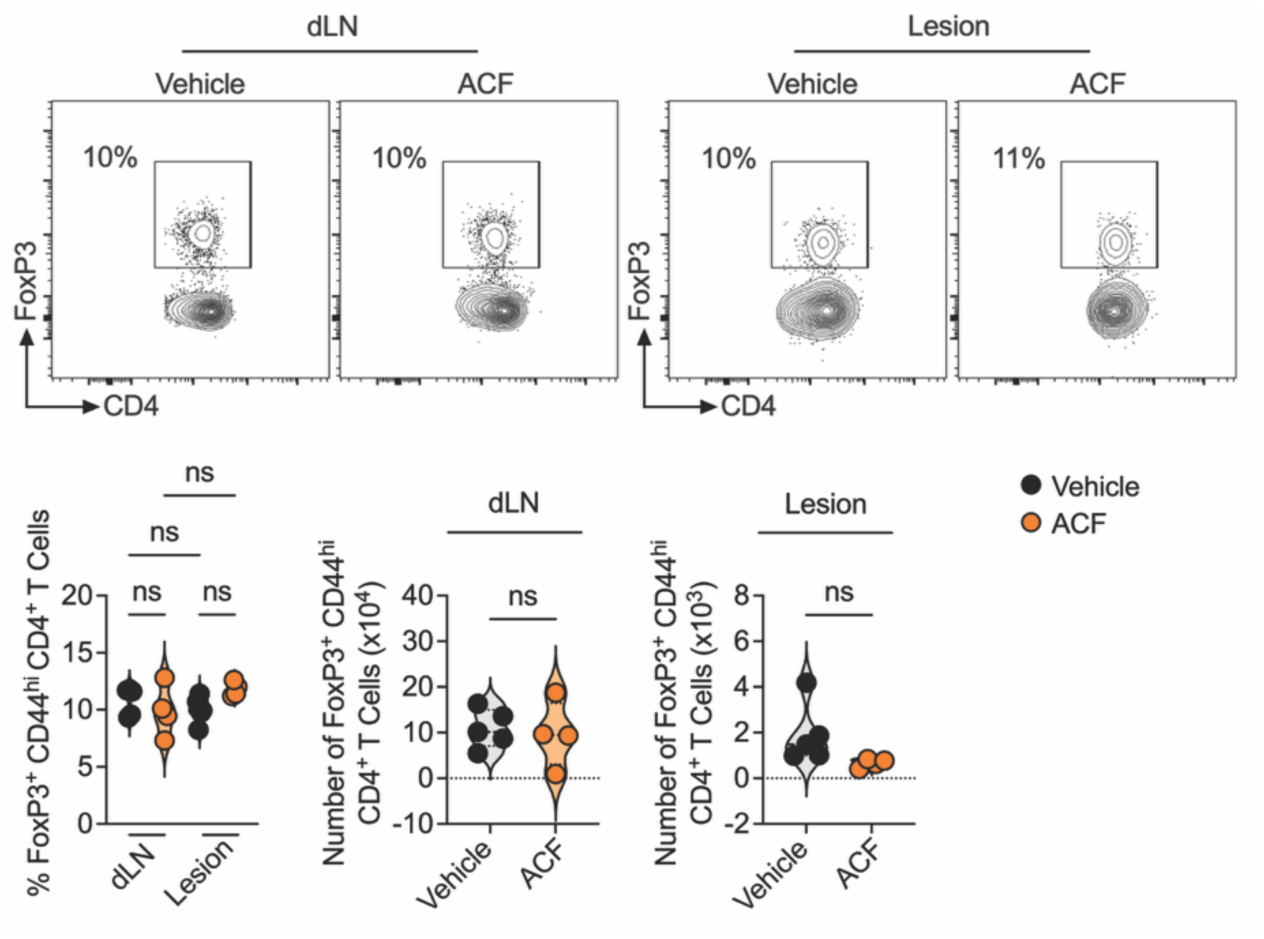
Treatment of *Leishmania*-infected mice with acriflavine does not impact T regulatory cells. C57BL/6 mice were infected with *Leishmania major* and treated daily with intraperitoneal injections of acriflavine or vehicle for 3 weeks. The frequency and number of FoxP3^+^ CD4 T cells in the dLN and lesion was analyzed by flow cytometry. Representative contour plot and scatter dot plots showing individual mice are representative of 2 experiments with at least 4 mice per experimental group. *P ≤ 0.05, ***P ≤ 0.001, and ****P ≤ 0.0001, by 2-tailed Student’s t-test (number) and 1-way ANOVA (frequency). Gating strategy: live, single cells, CD45, CD90, CD4, CD44^high^, FoxP3.

